# *Plasmodium falciparum* dipeptidyl aminopeptidase 3 activity is important for efficient erythrocyte invasion by the malaria parasite

**DOI:** 10.1101/202812

**Authors:** Christine Lehmann, Michele Ser Ying Tan, Laura E. de Vries, Ilaria Russo, Mateo Isidrio Sanchez, Dan E. Goldberg, Edgar Deu

**Affiliations:** Chemical Biology Approaches to Malaria Laboratory, The Francis Crick Institute, 1 Midland Road, London, United Kingdom; Radboud Institute for Molecular Life Sciences, Radboud University Nijmegen, Nijmegen, Netherlands; Faculty of Life Sciences, University of Manchester, Manchester, United Kingdom; Department of Genetic, Stanford School of Medicine, Stanford, United States.; Departments of Molecular Microbiology and Medicine, Washington University School of Medicine, St Louis, United States.

**Author notes:** Corresponding author: Edgar Deu.

## Abstract

Parasite egress from infected erythrocytes and invasion of new erythrocytes are essential for the exponential asexual replication of the malaria parasite, and both processes are regulated and mediated by proteases. The putative cysteine protease dipeptidyl aminopeptidase 3 (DPAP3) was previously suggested to be essential for parasite egress, but little is known about its biological function. Here, we demonstrate that DPAP3 has proteolytic activity, but contrary to previously studied DPAPs, removal of its prodomain is not required for activation. Interestingly, *P. falciparum* DPAP3 localizes to merozoite apical organelles from which it is secreted immediately before egress. Using a conditional knock out approach coupled to complementation studies with wild type or mutant DPAP3, we show that DPAP3 activity is critical for efficient RBC invasion and overall parasite replication, and demonstrate that it does not play a role in parasite egress. Overall, this study establishes DPAP3 as a key regulator of erythrocyte invasion.

---

Malaria is a devastating infectious disease caused by Apicomplexan parasites of the *Plasmodium* genus and is transmitted by Anopheles mosquitoes during a blood meal. After an initial asymptomatic liver infection, parasites are released into the blood stream where they replicate within red blood cells (RBCs). This asexual exponential growth is responsible for all the pathology associated with malaria, causing close to half a million deaths every year^1^. Over the last 15 years, the world has seen a significant decrease in malaria incidence mainly due to the distribution of insecticide-impregnated bed nets and the introduction of ACT (artemisinin-based combination therapy) as the standard of care for uncomplicated malaria^2^. However, the recent emergence of artemisinin resistance^3^ has made the identification of viable therapeutic targets extremely important^4,5^.

*P. falciparum* is the most virulent *Plasmodium* species accounting for most of malaria mortality. Its 48h asexual erythrocytic cycle consists of RBC invasion, intraerythrocytic parasite growth and division into 16-32 daughter merozoites, followed by parasite egress for further RBC invasion. Parasite egress and RBC invasion are key for parasite replication and blocking either one of these processes would lead to a quick drop in parasitemia and malaria pathology. Proteases have been shown to play essential roles in both processes and might therefore be viable therapeutic targets^6^.

RBC invasion is a multistep process involving initial recognition of RBC receptors by adhesin proteins on the surface of the merozoite (invasive extracellular parasite form), tight attachment to the RBC membrane (RBCM), reorientation of the merozoite apical end towards the RBCM, active invasion driven by an actin/myosin motor with invagination of the RBCM and formation of the parasitophorous vacuole (PV), and finally, sealing of the RBCM and PV membrane (PVM)^7,8^. The PV is a membrane-bound vacuole within which the parasite growths and replicates isolated from the RBC cytosol. Rupture of the PVM and RBCM is required for parasite egress and is mediated by proteases. In particular, subtilisin-like protease 1 (SUB1), an essential serine protease residing in apical secretory organelles known as exonemes, is released into the PV right before egress where it processes several proteins important for egress and invasion^9-13^. These include cleavage and likely activation of serine repeat antigen 6 (SERA6)^14^, an essential papain-fold cysteine protease^15,16^.

In a forward chemical genetic approach, *P. falciparum* dipeptidyl aminopeptidase 3 (DPAP3) was identified as a potential regulator of parasite egress acting upstream of SUB1^17^. DPAPs are papain-like cysteine proteases that cleave dipeptides off the N-terminal of protein substrates^18^. In that study^17^, the vinyl sulfone inhibitor SAK1 was shown to preferentially inhibit DPAP3 over other cysteine proteases. This compound arrests parasite egress at mid-micromolar concentrations, blocks processing of SUB1 substrates, and prevents proper expression and maturation of SUB1 and apical membrane antigen 1 (AMA1), a micronemal protein secreted onto the merozoite surface^9^ that is essential for RBC invasion^19^. These results led to the hypothesis that DPAP3 might act as a general maturase of secretory proteins involved in egress and invasion. However, while the function (or essentiality) of *P. falciparum* DPAP3 has not been validated genetically, in the rodent parasite *P. berghei*, DPAP3 knock out (KO) parasites are viable but replicate significantly slower^20-22^.

Here, we combine chemical, biochemical and conditional genetic approaches to show that DPAP3 is an active protease that resides in apical secretory organelles and that its activity is critical for efficient RBC invasion. We also provide very strong evidence showing that DPAP3 does not play a significant function in parasite egress.

## Results

### DPAP3 activity is important for parasite viability

Using single homologous recombination we were able to replace the endogenous catalytic domain of *dpap3* with a C-terminally tagged (GFP, mCherry or HA) version. However, our multiple attempts to replace the DPAP3 catalytic Cys with a Ser failed despite using the same homology region upstream of the catalytic domain (Figs. 1a-b and S1). Our attempts to KO *dpap3* by double homologous recombination also failed. These results strongly suggest that DPAP3 activity is critical for parasite replication. Parasites containing differently tagged *dpap3* were cloned by limited dilution, and the clones selected for this study will be referred to as DPAP3-GFP, DPAP3-mCh, and DPAP3-HA. Analysis of parasite extracts by western blot (WB) using a polyclonal antibody that targets the C-terminal half of DPAP3 showed a shift in migration pattern in accordance with the tag molecular weight (Fig. 1c).

**Figure 1.**
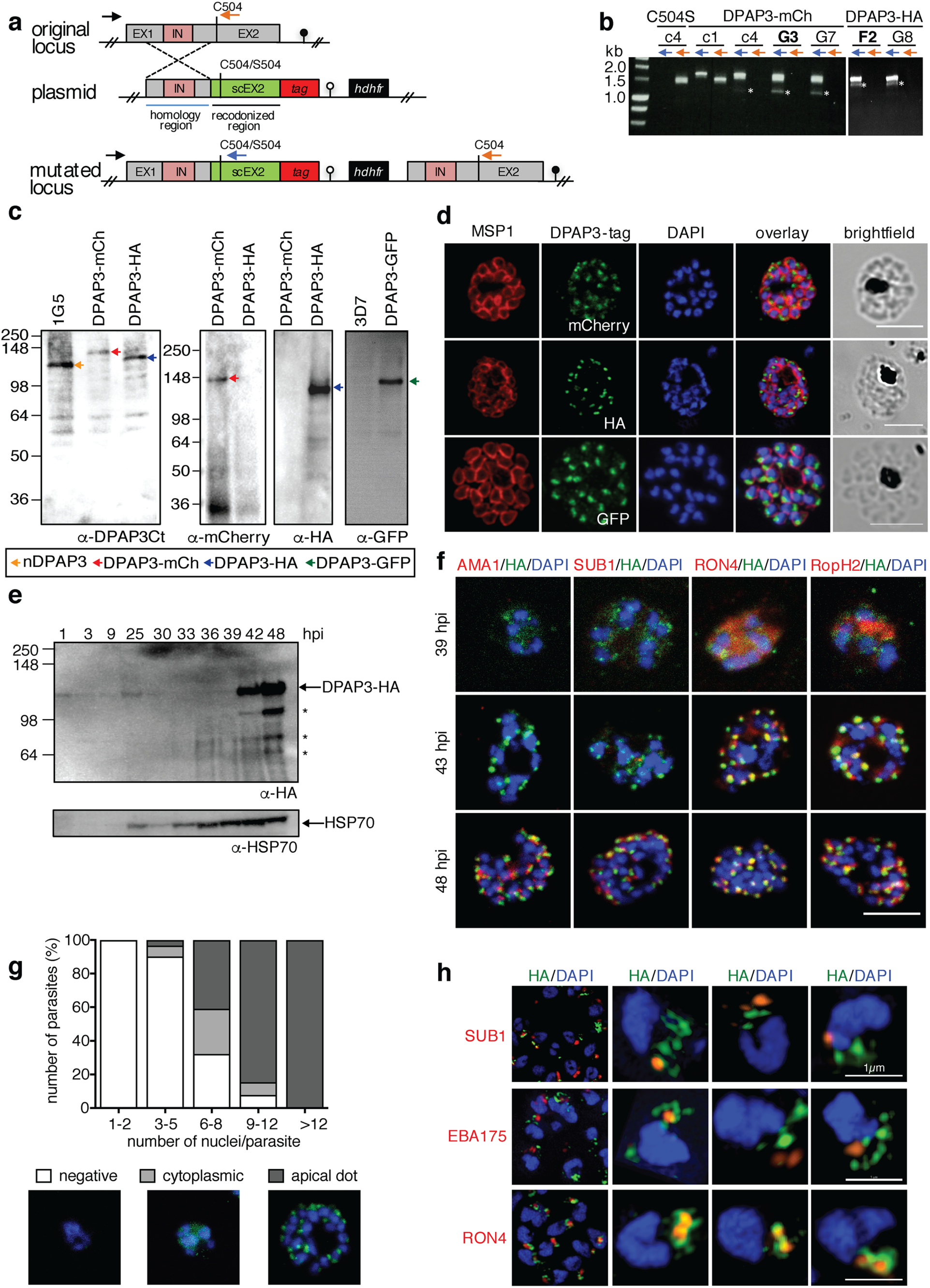
DPAP3 is expressed during schizogony and localizes at the apical pole. (**a**)Scheme showing the *dpap3* endogenous locus (exons, grey; intron, pink) targeted by single homologous recombination with plasmids containing a homology region (blue line), a recodonized C-terminal region (sc) tagged with HA3 or mCherry and containing either WT Cys504 or the C504S mutation, and the *hdhfr* drug resistant cassette. (**b**) Integration assessed by PCR on genomic DNA using the P21 (black arrow) and P22 (orange arrows) or P23 (blue arrow) primers for the endogenous or modified locus, respectively. Primer binding sites indicated in **a**. PCR obtained after one (c1) or four (c4) drug cycles, and for the DPAP3-mCh and DPAP3-HA clones are shown. Asterisks indicate non-specific PCR products. (**c**) WB analysis of schizont stage parasites from the DPAP3-mCh, -HA, and -GFP clones and the 1G5 and 3D7 control lines using anti-DPAP3-Ct, -HA, -mCherry, or -GFP antibodies. Bands corresponding to native or tagged DPAP3 are indicated with arrows. (**d**) IFAs of DPAP3-mCh, DPAP3-HA, and DPAP3-GFP mature schizonts stained with anti-MSP1 (red), and anti-mCherry, -HA, or - GFP (green) antibodies (confocal microscopy). (**e**) WB of DPAP3-HA parasite lysates harvested at different hours post RBC invasion (h.p.i). DPAP3-HA was detected using anti-HA antibody. HSP70 was used as a loading control. Asterisks indicate DPAP3 degradation products. (**f**) IFA of DPAP3-HA schizonts collected 39-48 h.p.i. and stained with anti-HA (green) and anti-AMA1 (microneme), anti-SUB1 (exoneme), anti-RON4 (rhoptry neck), or anti-RopH2 (rhoptry bulb) antibodies (all red). Obtained by confocal microscopy; single staining images are shown in Fig. S2. (**g**) Quantification of DPAP3-HA parasites showing negative, cytoplasmic or apical DPAP3 staining during schizogony. Samples were fixed from 36-48 h.p.i. and stained with anti-HA (green). Schizont maturity was assigned based on the number of nuclei. A representative image for each staining phenotype is shown. (**h**) IFA of C2-arrested DPAP3-HA schizonts analyzed by SIM. Parasites were stained with anti-HA (green) and anti-SUB1, anti-EBA175, or anti-RON4 antibodies (all red). Representative 3D-reconstruction images of merozoites are shown. In all IFAs DNA was stained with DAPI (blue); scale bars = 5µm except for merozoite images in **h** (1μm).

### PfDPAP3 localizes to an apical secretory organelle

In all our tagged lines, DPAP3 consistently localizes to the apical pole of merozoites (Fig. 1d). To determine when it is expressed, tightly synchronized DPAP3-HA and DPAP3-mCh parasites were collected every 3h throughout the erythrocytic cycle and were either lysed and analyzed by WB (Fig. 1e), or fixed for immunofluorescence analysis (IFA, Figs. 1f and S1c). Consistent with its transcription profile^23^, DPAP3 is most abundant in late schizonts and merozoites, but could also be detected in rings and trophozoites by WB. Although we could detect multiple processed forms of DPAP3 by WB, these processed forms are rarely observed in live parasites (Fig. 1c) and are likely an artefact of parasite lysis (see below).

By IFA, DPAP3 was first detected in young schizonts (6-8 nuclei) and seems to be expressed at the same time as rhoptry (RON4 and RopH2) and inner membrane complex (GAP45) proteins (Figs. 1f, S1 and S2). During schizont maturation, DPAP3 and rhoptry proteins localization changes from a diffuse and granular cytosolic staining to a clear punctuated apical staining in daughter merozoites (Figs. 1f-g, S1 and S2), probably reflecting protein trafficking and organelle biogenesis. By contrast, exonemal (SUB1) and micronemal proteins (AMA1) are expressed and localize to their apical organelles at a later stage (Figs. 1f and S2).

Using standard confocal microscopy, we could not observe consistent colocalization of DPAP3 with any apical organelle marker tested: AMA1 and EBA175 for micronemes, SUB1 for exonemes, RON4 and RopH2 for rhoptries, and Exp2 for dense granules (Figs. 1f and S2). We therefore decided to increase the resolution of our images by using structured illumination microscopy (SIM). By SIM we could clearly observe DPAP3 staining in small but well-defined dot-like structures at the apical pole of each merozoite in the DPAP3-HA (Figs. 1h and S3a) and DPAP3-GFP (Fig. S3b) lines. Although we did not observe colocalization with exoneme (SUB1), microneme (EBA175) or rhoptry (RON4, RopH2) protein markers (Figs. 1h and S3), DPAP3 seems to be closely associated with RON4, suggesting that it resides in apical organelles that surround the neck of the rhoptries (Fig. 1h).

### PfDPAP3 is secreted after PVM breakdown

Timely discharge of proteins from the different apical organelles is crucial to regulate parasite egress and RBC invasion^24^. Secretion of exonemal and micronemal proteins is mediated through activation of cGMP-dependent protein kinase G (PKG), which takes places 15-20 min before egress^9^. Release of SUB1 into the PV leads to parasite egress. Secretion of micronemal proteins, such as AMA1 or EBA175, onto the parasite surface is essential for merozoite attachment to the RBCM and invasion. To determine whether DPAP3 is secreted, we compared the level of DPAP3 present within parasites, in the PV and RBC cytosol (collected after saponin lysis), and in the culture supernatant at three different stages of egress: schizonts arrested before exoneme/microneme secretion with the PKG reversible inhibitor compound 2 (C2), schizonts arrested between PVM and RBCM breakdown using the general cysteine protease inhibitor E64, and free merozoites collected after egress. In E64-arrested schizonts, the erythrocyte is still intact but the RBCM is highly porated allowing leakage of RBC and PV proteins^25^. Parasite pellets, PV and RBC cytosol proteins, and proteins secreted in the culture supernatant under these three conditions were collected after treating the cultures with the fluorescent activity-based probe FY01^26^ in the presence or absence of the DPAP3 covalent inhibitor SAK1. FY01 is a cell-permeable probe that covalently modifies the catalytic Cys of DPAP3^17^. Fluorescently labelled DPAP3 can then be visualized by in-gel fluorescence in a SDS-PAGE gel. Pre-treatment of parasites with SAK1 prevents probe labelling resulting in the loss of the DPAP3 fluorescent band (Fig. 2a). Note that E64 does not inhibit DPAP3 since it does not block FY01 labelling. DPAP3 was detected in the culture supernatant and saponin fraction of E64-arrested or rupturing schizonts, but not in C2-arrested schizonts, indicating that it is secreted downstream of PKG activation but before merozoites become extracellular. The presence of DPAP3 in E64-arrested schizont pellets and free merozoites suggests only partial secretion before egress. DPAP3 secretion was confirmed by WB on the DPAP3-HA line and was only detected in the culture supernatant in the absence of C2 (Fig. 2b).

**Figure 2.**
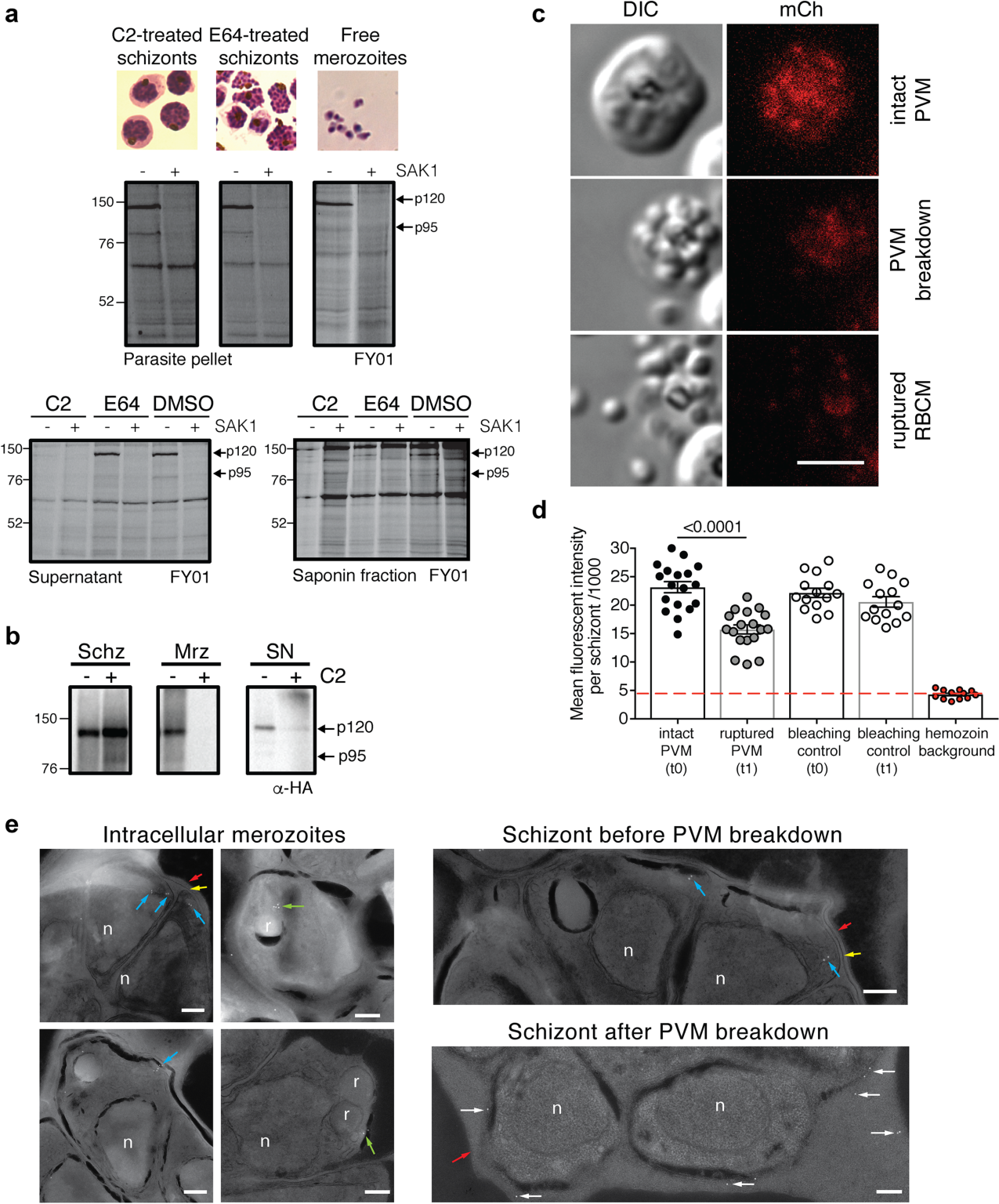
DPAP3 is secreted at the time of egress. (**a**) Rupturing (DMSO) and C2- or E64-arrested schizonts were labelled under intact conditions with FY01 in the presence or absence of the DPAP3 inhibitor SAK1. Parasite pellets from free merozoites and arrested schizonts (obtained after saponin lysis), proteins precipitated from the culture supernatant, and PV and RBC cytosol components (soluble saponin fraction), were run on a SDS-PAGE, and in-gel fluorescence measured in a flat-bed fluorescence scanner. Bands that disappear in the presence of SAK1 correspond to the p120 and p95 forms of DPAP3. The small proportion of p95 DPAP3 is likely produced after parasite lysis. (b) C2-arrested schizonts (DPAP3-HA) were either left on C2 or allowed to egress for 30min after C2 wash out. Unruptured schizonts, free merozoites, and proteins secreted into the culture supernatant were collected, and the presence of DPAP3-HA in each fraction visualized by WB using an anti-HA antibody. (**c**) Mature DPAP3-mCh schizonts were arrested with C2 for 3h, and egress observed by live video microscopy after C2 wash out (Video S1). The representative still-frame pictures show DIC and mCherry signal (red) before or after PVM breakdown, and after RBCM rupture. (**d**) Quantification of mCherry signal measured on consecutive frames before and after PVM breakdown. Around 20% of the signal originates from the hemozoin autofluorescence (red line). As a bleaching control, the mCherry signal of schizonts that did not egress was quantified at the corresponding time frames. (**e**) IEM section obtained from DPAP3-GFP parasites. Images of individual intracellular merozoites are shown on the left showing immunogold staining of DPAP3-GFP in close proximity to the rhoptries (green arrows) and at the apical end of merozoites (blue arrows). Images on the right show a representative section of schizonts before and after PVM (yellow arrow) breakdown. Staining of extracellular DPAP3-GFP (white arrows) was only observed in schizonts lacking a PVM. Rhoptries (r), nuclei (n), and the RBCM (red arrows) are indicated. Rabbit anti-GFP and colloidal gold-conjugated anti-rabbit antibodies were used. Bar graph = 200nm. IEM images obtained on the 3D7 control line are shown in Fig. S3d.

To determine the timing of DPAP3 secretion, DPAP3-mCh schizonts were arrested with C2 and parasite egress monitored by live microscopy after C2 wash out. Using this assay, PVM breakdown is clearly observable on the DIC channel when merozoites become more spread out within the RBC and their shape is better defined^12,25^. This is followed by RBCM rupture and merozoites dispersal (Fig. 2c and Videos S1a-c). Quantification of fluorescence signal in these egress video shows a 40% decrease in mCherry signal right after PVM breakdown but before RBCM rupture (Fig. 2d). Interestingly, conditional KO (cKO) of SUB1 has recently been shown to prevent breakdown of the PVM but not secretion of micronemal AMA1 onto the merozoite surface (Michael Blackman personal communication). This suggests that DPAP3 secretion occurs downstream of exoneme and microneme secretion, and might be triggered by PVM breakdown.

To confirm that DPAP3 is secreted into the RBC cytosol after PVM breakdown, we performed immunoelectron microscopy (IEM) on DPAP3-GFP schizonts collected at the time of egress (Fig. 2e). In schizonts containing an intact PVM, DPAP3-GFP staining was observed at the apical end of merozoites, and in some sections in close proximity to the rhoptries or enclosed within membrane bound vesicles, thus confirming our IFA results. In IEM images collected on schizonts after PVM breakdown, a high number of immuno staining was observed in the RBC cytosol. Rarely any DPAP3 staining was detected in the PV of schizonts with an intact PVM.

### Removal of the prodomain is not required for DPAP3 activation

DPAPs are composed of a signal peptide, an exclusion domain, a prodomain, and a C-terminal catalytic domain. Processing of DPAPs has mainly been studied for the mammalian homologue cathepsin C (CatC) and starts with removal of the signal peptide in the ER upstream of the exclusion domain N-terminal aspartate. This residue is critical for activity as it directly interacts with the free N-terminal amine of DPAP substrates^18^. Further processing consists of removal of the inhibitory prodomain and cleavage of the catalytic domain downstream of the catalytic Cys, which for CatC has been shown to be mediated by lysosomal cysteine cathepsins (Fig. 3a). Thus, each DPAP proteolytic unit is generally composed of three oligopeptides (exclusion domain and the N- and C-terminal portions of the catalytic domain), and CatC is composed of four proteolytic units^27^. Although *Plasmodium* DPAP1 seems to be processed in a similar way^28^, its final form is composed of a single proteolytic unit, thus showing that tetramerization is not required for DPAP activity (Fig. 3a).

**Figure 3.**
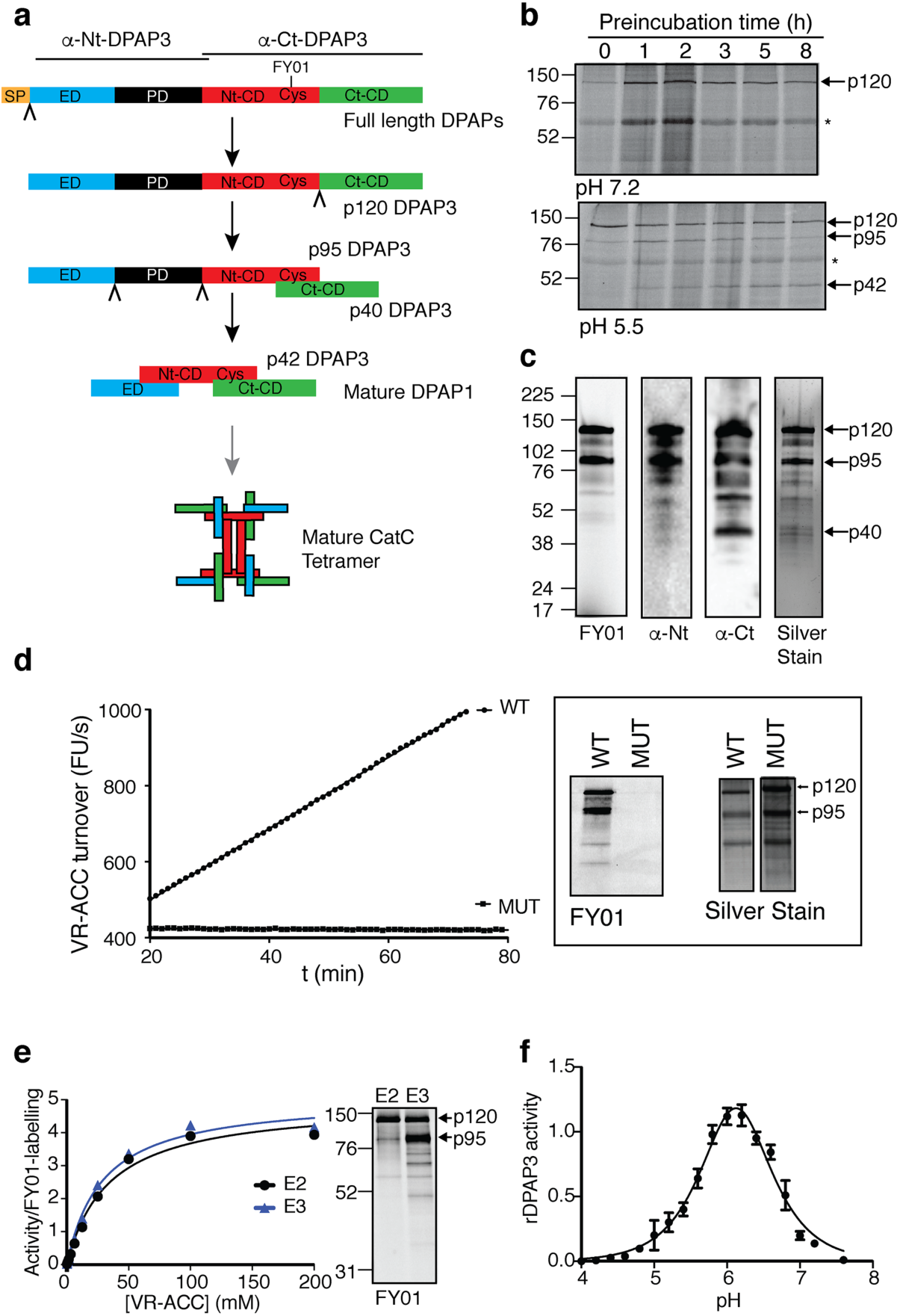
DPAP3 has proteolytic activity. (**a**) Schematic representation of DPAPs processing. The signal peptide (SP), exclusion domain (ED), prodomain (PD), and the N- and C-terminal portions of catalytic domain (Nt-CD and Ct-CD) are shown in different colours. Cleavage sited for removal of the SP and prodomain, and for processing of the catalytic domain are indicated by chevrons. A representation of fully processed monomeric DPAP1 and tetrameric CatC are shown. The predominant p120 DPAP3 active form is produced after removal of the signal peptide. We propose that the p95 DPAP3 active form is produced after cleavage of the catalytic domain. The regions recognized by anti-Nt-DPAP3 and anti-Ct-DPAP3 antibodies, and the position of the catalytic Cys covalently modified by FY01 are shown above the full-length protein scheme. (**b**) DPAP3 processing is an artefact of parasite lysis. Merozoite lysates were incubated for several hours under neutral (PBS, pH 7.2) or acidic (acetate buffer, pH 5.5) conditions before adding 1µM FY01 for 1h. Under acidic conditions, we observed a time dependent processing of the p120 form to the p95 and p42 forms. (**c**) Analysis of purified rDPAP3. Two main bands are detected by silver stain, both of which are strongly labelled by FY01 and recognized by the anti-Nt-DPAP3 and anti-Ct-DPAP3 antibodies. All other minor bands in the silver stain are also recognized by the DPAP3 antibodies and represent degradation products that could not be separated during purification. (**d**) Measurement of VR-ACC turnover and FY01 labelling for WT and C504S MUT recombinant DPAP3. Silver stain analysis shows equivalent amount of protein were obtain from the purification of WT and MUT DPAP3. (**e**) Michaelis Menten plots showing the VR-ACC turnover rate from two consecutive Ni-NTA elution fractions (E2 and E3) normalized to the total amount of FY01 labelling shown in the gel image. E2 contains predominantly the p120 form, and E3 a mixture of the p95 and p120 forms. (**f**) pH dependence of rDPAP3 activity measured at 10µM VR-ACC (n=3).

Three different isoforms of DPAP3 consistent with the canonical processing of DPAPs were previously shown to be labelled by FY01 in merozoite lysates under acidic conditions^17^ (p120, p95, and p42 forms). However, when intact schizonts or merozoites are directly boiled in SDS-PAGE loading buffer, we predominantly detect the p120 form either by WB or FY01 labelling (Figs. 1-2), suggesting that DPAP3 processing might be an artefact of parasite lysis. To confirm this, we show that incubation of merozoite lysates under acidic conditions results in a time-dependent processing of DPAP3, which is not observed at neutral pH (Fig. 3b).

Although FY01-labelling of the p120 ‘zymogen’ form indicates that its active site is accessible and contains a nucleophilic cysteine, it does not demonstrate that it can turnover substrates. To address this point we expressed full-length recombinant DPAP3 (rDPAP3) in insect cells using the baculovirus system^29^. Expression and purification of rDPAP3 from insect cells culture supernatant yielded predominantly the p120 and p95 forms (Fig. 3c), which are able to cleave the VR-ACC fluorogenic DPAP substrate^30^ (Fig. 3d). The detection of a 40kDa band by WB using an anti-DPAP3-Ct antibody suggests that cleavage of p120 DPAP3 into the p95 form occurs downstream of the catalytic Cys (Fig. 3c). However, given their molecular weights, both p120 and p95 forms must contain the prodomain, indicating that its removal is not required for DPAP3 activation. Finally, we show that mutation of the catalytic Cys to Ser completely inactivates DPAP3 (Fig. 3d). This inactive C504S mutant (MUT) was used as a negative control in our complementation studies (see below).

In one of our purifications we were able to separate the p120 form from a fraction containing a mixture of the p95 and p120 forms (Fig. 3e). Although rDPAP3 was more abundant in the latter fraction, when VR-ACC turnover was normalized to the total amount of FY01 labelling (i.e. p120+p95 labelling), both fractions showed identical *V*^max^ and *K*^m^ values, suggesting that these two forms are equally active, and that removal of the signal peptide is sufficient to activate DPAP3. Finally, we show that rDPAP3 is active under mild acidic conditions with an optimal pH of 6 (Fig. 3f). The facts that the p120 form of rDPAP3 is active, and that this is the predominant form found in live parasites, strongly suggest that this “zymogen” form is the one performing a biological function in parasites.

### Generation of DPAP3 cKO and complementation lines

Since we were unable to directly KO DPAP3, we generated DPAP3 cKO lines on the 1G5 parasite line background that endogenously expressed DiCre^31^. In the DiCre system, Cre recombinase is split into two domains fused to rapamycin (RAP) binding domains. Addition of RAP triggers dimerization and activation of DiCre, leading to rapid recombination of specific DNA sequences known as *LoxP* sites^32,33^. We used this system to conditionally truncate the catalytic domain of DPAP3 rather than excising the full gene to prevent potential episomal expression of DPAP3 after excision.

Two independent strategies were used (Fig. 4a), which differ in how the first *LoxP* site was introduced within the *dpap3* open reading frame (ORF). In both cases, the second *LoxP* site was introduced downstream of the 3’-UTR. The first one was inserted either within an Asn-rich region of DPAP3 predicted not to interfere with folding or catalysis, or within an artificial intron (*LoxPint*), which does not alter the ORF of the targeted gene and that has been shown to be well tolerated in several *P. falciparum* genes^12,34,35^. In both instances, the recodonized catalytic domain was tagged with mCherry such that RAP-induced truncation would result in the loss of mCherry signal. A control line containing only the 3’-UTR *LoxP* site was also generated. After transfection of 1G5 parasites, drug selection, and cloning, three DPAP3cKO clones (F3cKO and F8cKO with *loxPint*, and A1cKO with *loxP* in Asn-stretch), and the E7ctr line (only one *loxP*) were selected for further studies. Evidence of integration by PCR is shown on Fig. S4b. Analysis of genomic DNA of DMSO- or RAP-treated cKO lines by PCR showed highly efficient excision (Fig. 4b) resulting in the loss of DPAP3-mCh expression in mature schizonts (Fig. 4c-d). Although we consistently achieved more than 95-99% excision efficiency (Fig. 5a), a fraction of non-excised parasites was always present after RAP treatment, which explains the presence of a non-excision DNA band by PCR after RAP treatment (Fig. 4b).

**Figure 4.**
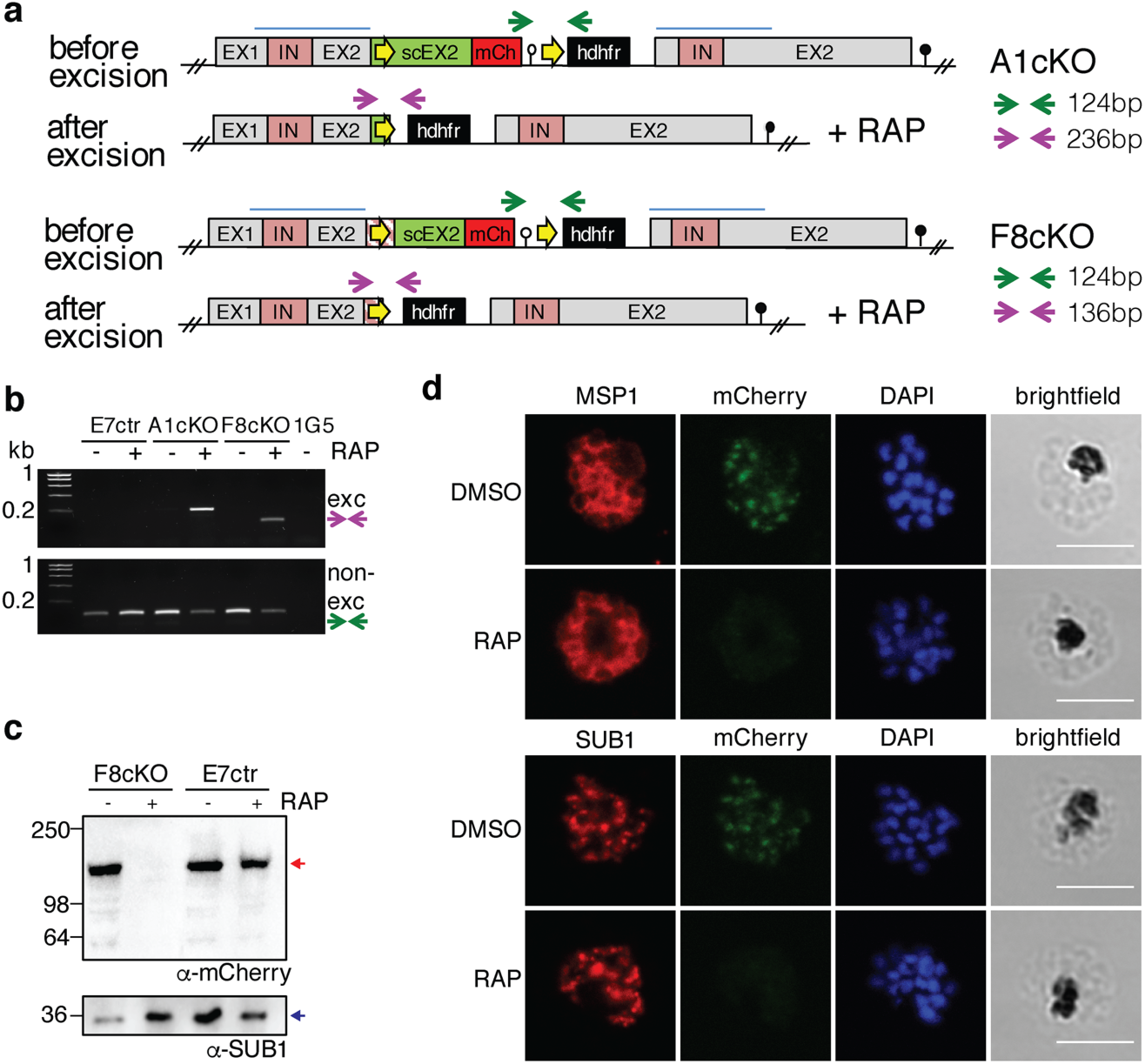
Generation of DPAP3 cKO Lines. (**a**)Schematic representation and assessment of the *dpap3* recombinant genetic locus before and after RAP-mediated excision for the F8cKO and A1cKO lines. Wild type exons (EX) and intron (IN) sequences are depicted with grey and pink boxes, respectively. The homology regions used for single crossover recombination are indicated with blue lines, and the recodonized 3’ end of the second exon is shown in green (scEX2). *LoxP* sites (yellow arrows) were introduced downstream of the *P. berghei* 3’UTR (white circle) and either within the ORF of scEX2 (A1cKO) or upstream of scEX2 (F8cKO) as a *LoxPint* artificial intron (pink striped box). The *mCherry* coding sequence (red box) and the *hdhfr* resistance cassette (black box), and the displaced endogenous *dpap3* locus with its 3’UTR (black circle) are also shown. Arrows indicate primers annealing sites used for diagnostic PCR of excised (purple) and non-excised (green) loci. (**b**) Diagnostic PCR showing excision at the *dpap3*-locus. PCR was performed on genomic DNA collected from the E7ctr, A1cKO and F8cKO lines 24h after DMSO or RAP treatment. Genomic DNA from the parental 1G5 line was used as a negative control. Excision and non-excision PCR products are indicated with purple and green arrows, respectively. Excision product was only observed after RAP treatment of the cKO lines. The presence of a non-excised PCR product after RAP treatment indicate that excision is not 100% efficient. (**c**) WB analysis showing highly efficient loss of DPAP3 upon RAP treatment. Schizonts collected 45h after DMSO or RAP treatment of E7ctr and F8cKO parasites were saponin lysed, and the parasite pellet analyzed by WB using an anti-mCherry antibody (red arrow). SUB1 was used as a loading control. (**d**) IFA analysis of mature schizonts showing the loss of DPAP3 signal after RAP treatment. Ring-stage F8cKO parasites were treated with DMSO or RAP for 3h and fixed for IFA analysis at 48 h.p.i. Slides were stained with anti-mCherry (green) and anti-MSP1 (red) or anti-SUB1 (red) antibodies. DNA was stained with DAPI (blue); scale bar: 5µm.

**Figure 5.**
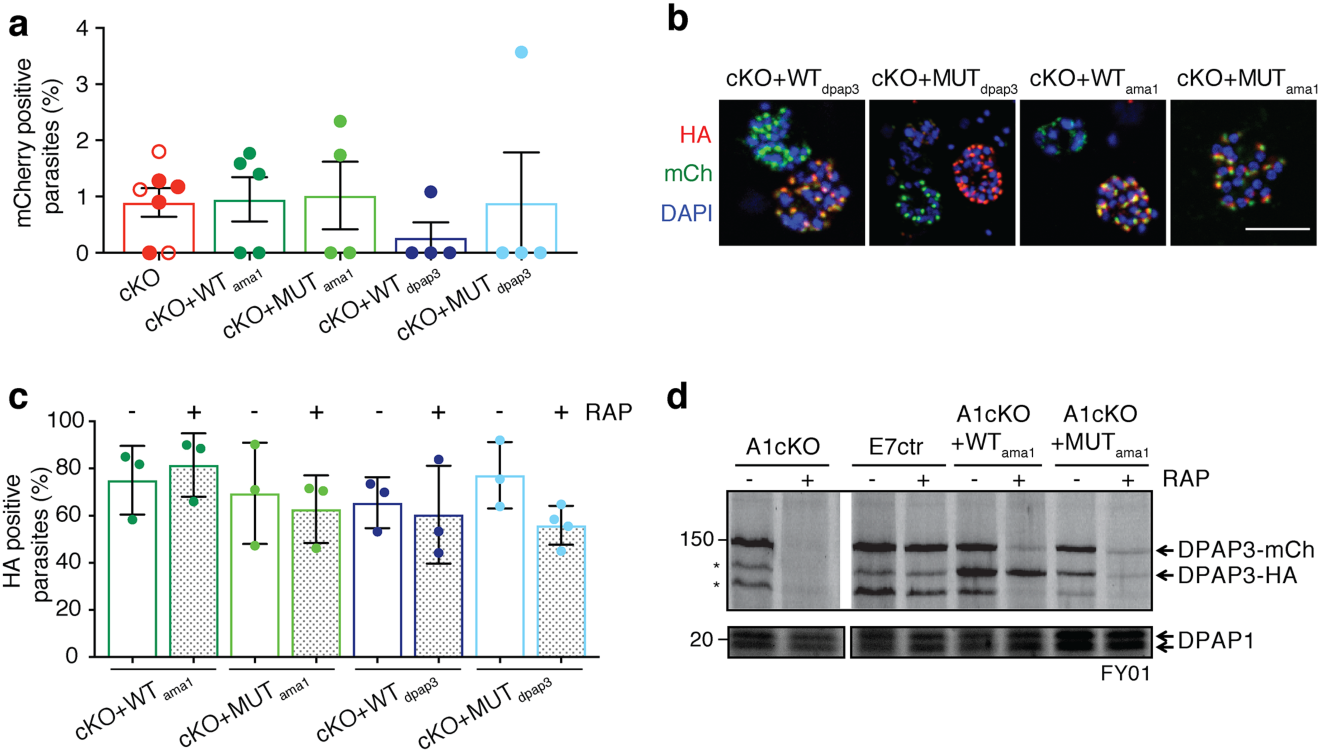
Generation of complementation lines. (**a**)Quantification of excision efficiency for DPAP3cKO and complementation lines. Schizonts collected after DMSO or RAP treatment were stained with anti-mCherry and anti-SUB1 antibodies. SUB1 staining was used as a marker of schizont maturity. The amount of mCherry positive schizonts was quantified in relation to the total amount of mature schizonts. In DMSO treated parasites, all mature schizonts were mCherry positive (100%). The amount of non-excised parasites after RAP treatment was <5% in all biological replicates for all the cKO and complementation lines tested. Each circle corresponds to a different biological replicate (>100 schizonts were analyzed per biological isolate). Filled circles correspond to F8cKO and its complementation lines, and empty ones to the A1cKO line. (**b**) IFA analysis of the complementation lines showing colocalization of chromosomal DPAP3-mCh and episomal DPAP3-HA expressed under that *dpap3* or *ama1* promoters. Mature schizonts from the F8cKO+WT_dpap3_, F8cKO+MUT_dpap3_, F8cKO+WT_ama1_, and F8cKO+MUT_ama1_ were fixed and stained with anti-HA (red) and anti-mCherry (green). DNA was stained with DAPI (blue); scale bar: 5µm. Single coloured images are shown in Fig. S4c. (**c**) Quantification of the amount of DPAP3-HA positive schizonts in the complementation lines. Schizonts were fixed at 48 h.p.i. and stained with anti-HA and anti-SUB1 antibodies. Only 60-80% of mature schizonts show positive HA staining with no difference between DMSO and RAP treated parasites. More than 100 schizonts per biological replicate were analyzed. (**d**) FY01 labelling of cKO and complementation lines. After DMSO or RAP treatment at ring stage, C2-arrested schizonts were collected, lysed, and labelled with FY01. Labelling of chromosomal DPAP3-mCh is clearly visible as a band around 150kDa along with some post-lysis degradation products indicated by asterisks. The loss of this 150kDa upon RAP treatment is observed in all lines except the E7ctr. Episomal WT DPAP3-HA is labelled by FY01 independently of RAP treatment and co-migrates with one of the degradation products of DPAP3-mCh at 125kDa. No labelling of MUT DPAP3-HA was observed. DPAP1 labelling by FY01 is shown as a loading control.

To confirm that any phenotypic effect observed upon conditional truncation of DPAP3 is due to the loss of DPAP3 activity, A1cKO and F8cKO parasites were transfected with plasmid expressing WT or MUT DPAP3-HA under the control of the *dpap3* or *ama1* promoters (Fig. S4a), resulting in the following complementation lines: F8cKO+WT_dpap3_, F8cKO+MUT_dpap3_, F8cKO+WT_ama1_, F8cKO+MUT_ama1_, A1cKO+WT_dpap3_, A1cKO+WT_ama1_, A1cKO+MUT_ama1_. All complementation lines grew normally before RAP treatment and showed no apparent delay in parasite development.

IFA analysis confirmed colocalization between chromosomal DPAP3-mCh and episomal DPAP3-HA (Fig. 5b). Episomal expression was only high enough to be detected by IFA in 60-80% of schizonts, but was independent of RAP treatment. This is probably due to different levels of episomal expression and plasmid segregation in schizonts (Fig. 5c). Efficient, but not complete, truncation of DPAP3-mCh was observed in all our complementation lines (Fig. 5a). Labelling of DPAP3 with FY01 in parasite lysates from these lines show clear labelling of chromosomal DPAP3-mCh and episomal WT DPAP3-HA but not MUT DPAP3-HA (Fig. 5d). As expected, RAP treatment results in a decrease of labelling of chromosomal DPAP3-mCh but not of episomal WT DPAP3-HA.

### Conditional knock out of DPAP3 reduces viability of blood stage parasites

To measure the effect of DPAP3 truncation on parasite replication, we used the recently published plaque assay^16^ where the wells of a 96-well flat bottom plate containing a thin layer of blood were seeded with ~10 iRBCs/well. After 10-14 days, microscopic plaques resulting from RBC lysis can be detected. RAP treatment of our cKO lines resulted in 90% less plaques, an effect that could be partially rescued through episomal complementation with WT but not MUT DPAP3 (Fig. 6a, Table S1). After RAP treatment of the F3cKO and F8cKO lines, some wells contained a single plaque, suggesting that only one clonal parasite population grew in that well. Parasites present in 12 of these wells were propagated and *dpap3* excision checked by PCR. All contained non-excised parasites but *dpap3* excision was detected in three cultures (Fig. 6b). The presence of excised parasites in some of the wells suggests that DPAP3KO parasites replicate less efficiently and might not have had enough time to form a microscopic plaque within the 14 days of the assay. That said, the presence of non-excised parasites in all samples indicates that WT parasites quickly outcompeted DPAP3KO ones.

**Figure 6.**
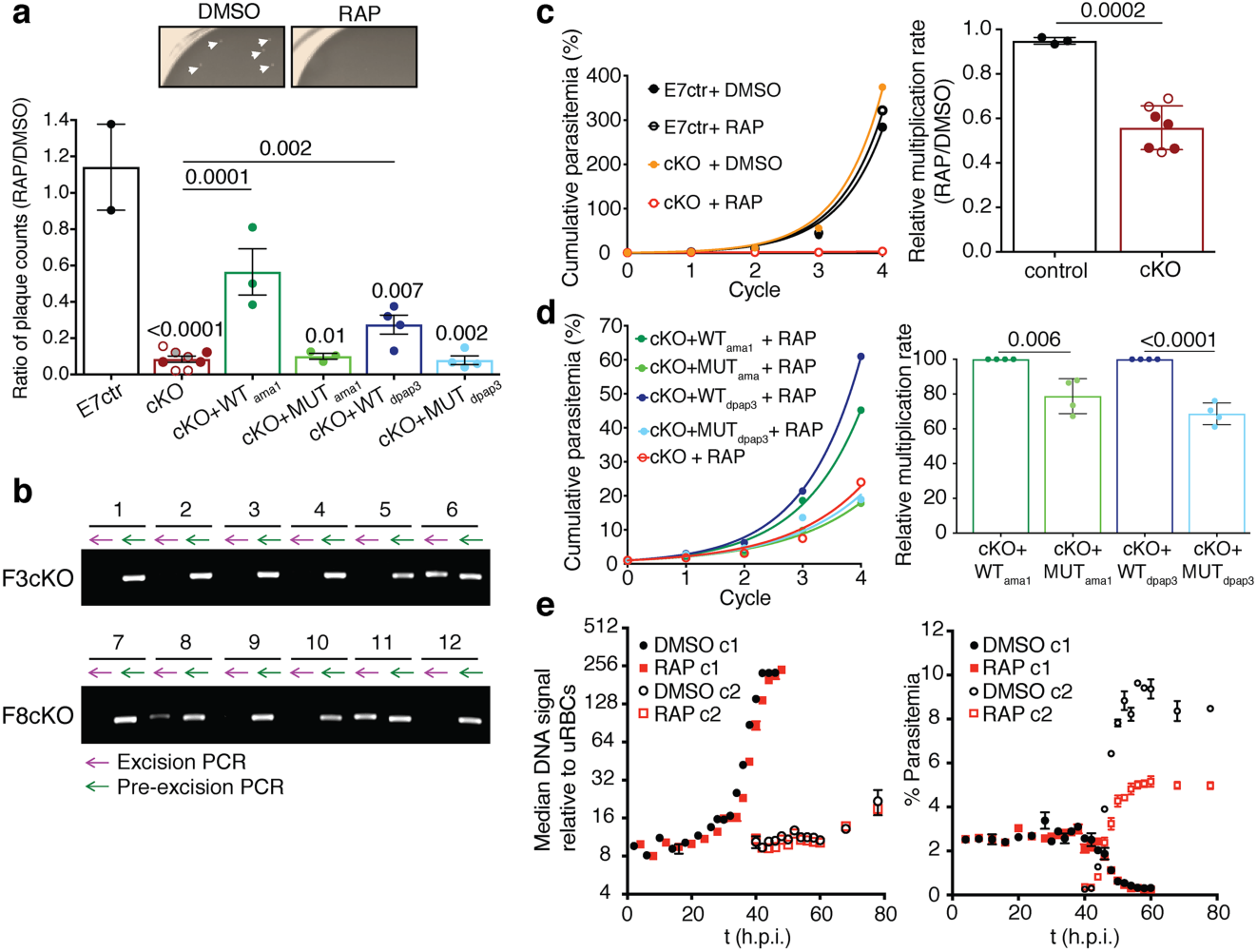
Conditional knock-out of DPAP3 leads to a severe growth defect. (**a**) Evaluation of parasite growth by plaque assay. Our different parasite lines were treated with DMSO or RAP and grown in flat bottom 96-well plates at 10 iRBC/well, and the number of plaques counted after 10-14 days (Table S1). The ratio between RAP and DMSO treated parasites is plotted in the bar graph. Representative images of two wells from DMSO or RAP treatment of F8cKO parasites are shown (plaques are indicated by arrows). (**b**) Plaques originating from RAP-treatment of cKO lines contain non-excised parasites. Parasites from 12 wells that only contained a single plaque after RAP treatment of F3cKO and F8cKO parasites were expanded, and their genomic DNA analyzed for excision (purple arrow) and non-excision (green arrow) by PCR. (**c-d**) Effect of DPAP3 cKO on parasite replication. The growth of DMSO- or RAP-treated E7ctr and DPAP3cKO lines was monitored by FACS for four cycles. Cumulative percentage parasitemia was fitted to an exponential growth model. Representative growth curves for F8cKO and E7ctr parasites are shown. The bar graph shows the effect of RAP treatment on the culture multiplication rate relative to DMSO treatment. (**d**) Representative replication curves obtained after RAP treatment of F8cKO and its complementation lines. Bar graph compares the multiplication rate after RAP treatment of DPAP3cKO lines complemented with WT or MUT DPAP3. (**a**, **c** and **d**) Each circle indicates a biological replicate: filled, F8cKO and its complementation lines; empty, A1cKO; grey, F3cKO. Error bars represent standard deviations. Only student’s t test significant values are shown. (**e**) Effect of DPAP3 KO on parasite development. Tightly synchronized A1cKO parasites pre-treated with DMSO or RAP were monitored over 76h by FACS based on DNA content (Hoechst staining). <u>Left Graph:</u> No differences in DNA content was observed between DMSO and RAP treatment. <u>Right</u> <u>Graph:</u> The time-dependent decrease of parasites belonging to the 1^st^ cycle (c1) after DMSO or RAP treatment coincides with an increase of a parasite population belonging to 2^nd^ cycle (c2), thus effectively monitoring egress and invasion. Results are the mean ± standard deviation of three technical replicates.

To test this hypothesis, we perform standard parasite replication assays after RAP or DMSO treatment. To prevent parasite overgrowth, cultures were diluted 10-fold in fresh blood and media whenever parasitemia reached 5%. RAP treatment of DPAP3cKO parasites results in a 10- to 15-fold decrease in parasitemia after 3-4 cycles, corresponding to an overall 50% decrease in multiplication rate per cycle compared to DMSO treatment (Fig. 6c). However, these values underestimated the importance of DPAP3 on parasite replication since 5 cycles after RAP treatment, 60% of parasites are non-excised and express DPAP3-mCh (Fig. S5a), thus proving that the small fraction of non-excised parasites quickly outcompete the excised ones. After RAP treatment, parasites complemented with WT DPAP3 grew significantly faster than those complemented with MUT DPAP3 (Fig. 6d). Importantly, our multiple attempts to clone DPAP3KO parasites after RAP treatment failed, indicating that DPAP3 activity is required for parasite propagation under our culturing conditions.

To determine which point of the erythrocytic cycle is disrupted by the loss of DPAP3, a tightly synchronized culture of A1cKO parasites at ring stage was treated with DMSO or RAP for 3h, and the culture was monitored for the following 80h. Samples were collected every 2-4h, stained with Hoechst, and analyzed by FACS. No significant difference in DNA staining was observed between WT and KO parasites, suggesting that DPAP3 is not required for intracellular development (Fig. 6e, left graph). Quantification of iRBCs belonging to the first or second cycle after treatment shows that DPAP3KO parasites egress at the same time as WT but produce **~**50% less rings (Fig. 6e, right graph). This suggests that DPAP3 is only important for RBC invasion, which is in direct contradiction with its previously suggested role in egress^17^.

### PfDPAP3 does not play a role in parasite egress

Despite being expressed early during schizogony (Fig. 1f), we did not observe any delay between 36-48 h.p.i. (Fig. 6e). This was confirmed by IFA by counting the number of mature schizonts at the end of the cycle after excision. Apical localization of SUB1 to the exonemes was used as a marker for schizont maturity. No significant difference was observed between DMSO and RAP treatment of cKO or complementation lines (Fig. S5b). Importantly, proper localization of MSP1, SUB1, AMA1, EBA175, RON4, and RopH2 was observed in all RAP-treated cKO lines (Figs. 4d and S5c-d).

Previously published work using the SAK1 inhibitor showed arrest of egress upstream of SUB1 activation^17^. The data suggested that DPAP3 might be important for parasite egress. Although we have been able to reproduce these results, we show that SAK1 treatment of schizonts 6h before egress arrests schizogony upstream of SUB1 and AMA1 expression rather than parasite egress (Fig. S6). This explains why no SUB1 or AMA1 could be detected by WB in the previous study^17^. In addition to SAK1, we also synthesized a more selective inhibitor by replacing the nitro-tyrosine N-terminal residue of SAK1 with *L-*Trp (*L*-WSAK) and its diastereomer negative control containing *D-*Trp (*D-*WSAK). To test the specificity of these inhibitors, merozoite or schizont lysates were pre-incubated with a dose response of compound followed by 1h labelling with FY01 (Fig. 7a). Although SAK1 blocks labelling of DPAP3 at lower concentration than *L-*WSAK, it also inhibits all the falcipains (FP1, FP2, and FP3) above 5μM, while *L-*WSAK only inhibits other targets above 200μM. As expected, the *D-* WSAK control is unable to inhibit any of the labelled cysteine proteases and is at least 100-fold less potent than *L-*WSAK at inhibiting DPAP3 (Fig. 7a). We then compared the effect of these inhibitors in parasite egress on the A1cKO line. Surprisingly, all blocked egress independently of RAP treatment (Fig. 7b), and no difference in potency between *L-* and *D-* WSAK was observed. These results prove that these vinyl sulfone compounds do not act through inhibition of DPAP3 but rather through off-target or toxicity effects.

**Figure 7.**
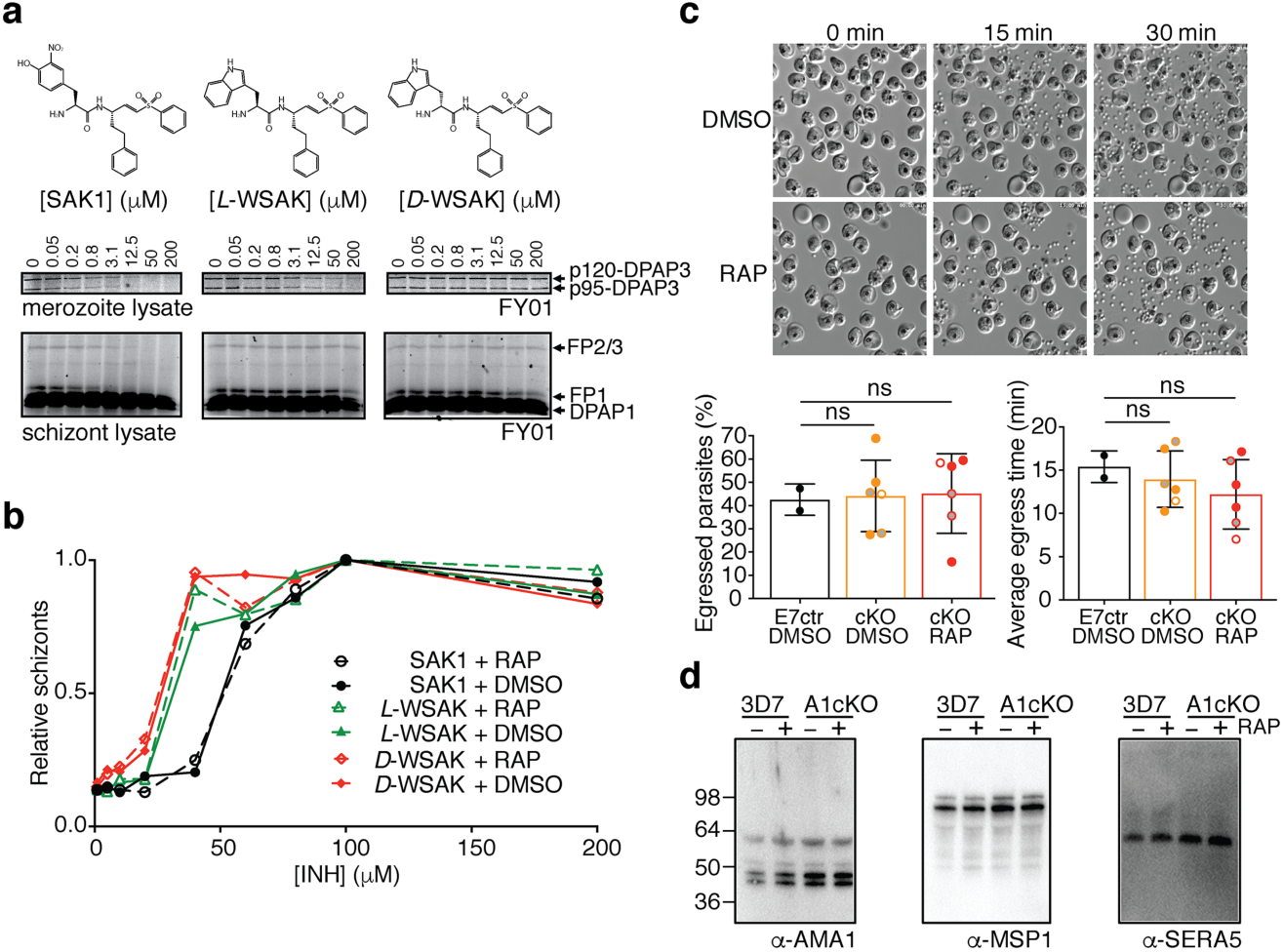
Role of DPAP3 in parasite egress. (**a**) *L*-WSAK is a more selective DPAP3 inhibitor than SAK1. The structures of SAK1, *L-*WSAK and *D*-WSAK are shown. Merozoite or schizont lysates were pre-incubated with a dose response of inhibitor for 30min followed by FY01 labelling. Samples were run on an SDS-PAGE gel, and the gel scanned on a flatbed fluorescence scanner. Bands corresponding to each of the labelled cysteine proteases are indicated by arrows. (**b**) Effect of inhibitors on egress. DMSO or RAP treated A1cKO parasites were treated at schizont stage with a dose response of inhibitor for 24h. The accumulation of schizonts upon inhibitor treatment was quantified by FACS. (**c**) Analysis of egress by video microscopy. C2-arrested schizonts obtained from DMSO- or RAP-treated of A1cKO, F8cKO, F3cKO or E7ctr parasite lines were monitored by time-lapse DIC microscopy for 30min after C2 washout. Representative still images taken at 0, 15, and 30min are shown for F8cKO parasites. The full time-lapse video can be seen in Video S2. The percentage of schizonts that egressed during this 30min time-lapse (left graph) and the time at which each individual schizont ruptured (right graph) are shown. Bar graphs show mean values ± standard deviation; circles show individual biological replicates (filled for F8cKO, empty for A1cKO, and grey for F3cKO). (**d**) WB analysis of culture supernatant collected after egress of 3D7 and A1cKO after DMSO or RAP treatment. No differences in the processing of AMA1, MSP1 or SERA5 was observed as a result of DPAP3 truncation.

As a final proof to show that DPAP3 is not involved in parasite egress, we arrested DMSO or RAP treated F8cKO schizonts with C2 and monitored egress by live microscopy after removal of the PKG inhibitor. Analysis of these videos showed no significant difference in the number of schizonts that ruptured, nor on how fast merozoites egressed after C2 wash out (Fig. 7c and Video S2). Also, we could not detect differences in the levels and/or processing of AMA1^36^ or SUB1 substrates (SERA5^10^ and MSP1^13^) between DMSO and RAP treated parasites (Fig. 7d). These findings together with the lack of colocalization between DPAP3 and SUB1 (Figs. 1h and S3) clearly demonstrate that DPAP3 is not responsible for proper processing and activation of SUB1, and that DPAP3 does not play a role in egress. However, its localization in an apical secretory organelle (Figs. 1-2) and the time-course analysis of the A1cKO line (Fig. 6e) strongly suggest a function in RBC invasion.

### PfDPAP3 is important for efficient attachment of merozoites to the surface of RBCs

Mature schizonts obtained after DMSO or RAP treatment of our different parasite lines were incubated with fresh RBCs for 8-14 h, fixed, and the population of schizonts and rings quantified by FACS (Fig. 8a). On average, we observed a 50% reduction in the number of infected erythrocytes after RAP treatment of our cKO lines. This invasion defect could be rescued by episomal expression of WT but not MUT DPAP3 independently of the promoter used (*ama1* or *dpap3*), indicating that DPAP3 activity is important for RBC invasion (Fig. 8b).

To determine which invasion step is impaired by the loss of DPAP3, DMSO- or RAP-treated cKO parasites were arrested at schizont stage with C2, incubated with fresh RBCs after C2 washout, and samples collected at different time points for FACS analysis. Fluorescent wheat germ agglutinin (WGA-Alexa647) binds to lectins on the RBC surface and when combined with Hoechst staining allows us to differentiate free merozoites from iRBCs (Fig. 8c). Quantification of the different parasite stage populations over time clearly shows a decrease in the number of rings upon RAP treatment, with an inversely proportional increase in the number of free merozoites (Figs. 8c and S7a).

The RBC population showing similar levels of DNA signal as free merozoites is mainly composed of ring stage parasites but likely contains a proportion of extracellular merozoites attached to the surface of the erythrocyte. These two populations were differentiated by staining the samples with the monoclonal MSP1 antibody m89.1, whose epitope is within the portion of MSP1 that is shed during invasion (Fig. 8d). Using this assay we did not observe a significant difference in the number of attached merozoites relative to the number of rings between DMSO and RAP treatment of cKO lines. This suggests that after DPAP3KO parasites tightly attach to the RBC surface they invade as efficiently as WT parasites, and therefore that DPAP3 is likely important for the initial recognition of the RBC.

**Figure 8.**
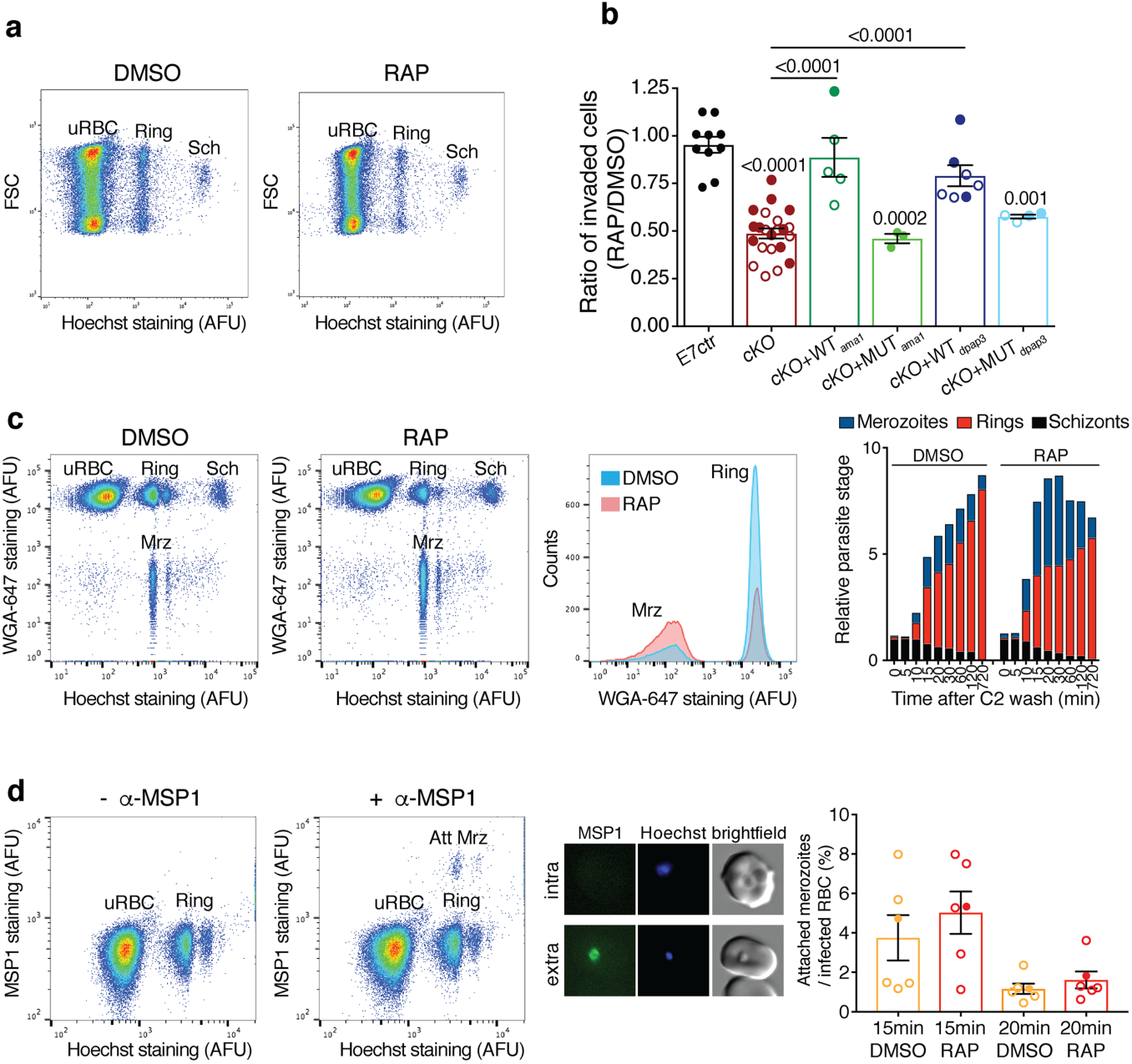
Role of DPAP3 in RBC invasion. (**a**) Representative FACS plot (forward light scattering vs. Hoechst staining) showing a decrease in invasion of the A1cKO upon RAP treatment. The populations of uRBCs, rings, and schizonts (Sch) are indicated. (**b**) Analysis of invasion efficiency of DPAP3 cKO and complementation lines. Schizonts collected 45h after DMSO or RAP treatment were incubated with fresh erythrocytes for 8-14h, fixed, stained with Hoechst, and analyzed by FACS. Shown is the ratio in invasion efficiency between RAP- and DMSO-treated parasites. Filled and empty circles represent individual biological replicates for the F8cKO and A1cKO and their corresponding complementation lines, respectively. Student’s t test significance values between cell lines are shown above the lines, or above each bar when comparing to the E7ctr. Only significant p-values are shown. (**c**) FACS analysis of extracellular merozoites. C2-arrested A1cKO schizonts pretreated with DMSO or RAP were incubated with fresh RBCs after C2 removal. Samples were collected at the indicated time points, fixed, and stained with Hoechst and WGA-Alexa647. The FACS plot and histogram show samples collected 20min after C2 washout. Free merozoites (Mrz) show positive staining for DNA but negative for WGA-Alexa647. Quantification of the different parasite stage populations over time is shown on the bar graph; biological replicates are shown in Fig. S7. (**d**) Quantification of attached merozoites by flow cytometry. Samples collected 15 and 20min after C2 washout during invasion assays (performed as in c), were stained with Hoechst and anti-MSP1 antibody (anti-mouse Alexa488 as secondary antibody). Because MSP1 is shed during invasion, merozoites attached to the RBCM (Att Mrz) can be differentiated from intracellular parasites as the cell population positive for DNA and MSP1 staining. FACS plots comparing anti-mouse Alexa488 staining in samples treated with or without the anti-MSP1 antibody. MSP1 staining (green) of attached merozoites under these conditions was confirmed by microscopy (central panel). Quantification of the population of attached merozoites relative to the ring population is shown on the bar graph. Circles represent different biological replicates: filled for F8cKO and empty for A1cKO. No significant difference was observed between DMSO and RAP treatment.

## Discussion

This study provides the first characterization of the biological function of DPAP3 in parasite development. We have shown that DPAP3 is critical for efficient RBC invasion and parasite replication, that it is expressed early during schizogony and localizes to small apical organelles that are closely associated with the neck of the rhoptries, and that it is secreted after PVM breakdown but before parasite egress. We have also demonstrated that DPAP3 has dipeptidyl aminopeptidase activity but contrary to other DPAPs, removal of the prodomain is not required for activation. Our cKO and complementation studies provide strong evidence that DPAP3 activity is only required for efficient RBC invasion, but not for intracellular parasite development or parasite egress. Finally, our initial characterization of the invasion defect associated with the loss of DPAP3 suggests that this protease plays an important role in the initial attachment of merozoites to the RBC surface.

The fact that DPAP3 is expressed early during schizogony, that it localizes in secretory organelles, and that it is active under mild acidic conditions, suggest that DPAP3 might process its substrates before it is secreted. It is difficult to speculate about the nature of these substrates since we could not colocalize DPAP3 with any of the tested apical markers. That said, DPAP3 might process transmembrane or GPI-anchored substrates residing within the same apical organelle but relocalizing to the merozoite surface after secretion (similarly to micronemal and some rhoptry proteins^8^). Alternatively, DPAP3 might reside in an apical late endosome compartment and might be important for sorting secretory proteins to their correct apical organelles by selectively trimming their N-termini after signal peptide removal. Indeed, the N-terminal sequence downstream of the signal peptide has been shown to be important for proper localization of rhoptry and micronemal proteins both in *P. falciparum*^37,38^ and *T. gondii*^39,40^. Micronemes are trafficked to the merozoites apex underneath the IMC before releasing their protein content, thus coming in very close proximity to the neck of the rhoptries^41,42^. DPAP3 might get in contact with proteins from other secretory organelles right at the time of secretion, similarly to how micronemal CyRPA and PfRipr come together with rhoptry PfRh5 at the merozoite apex after egress^43^. At this stage, we cannot discard the possibilities that DPAP3 might process its substrates after secretion or during trafficking (in the ER or Golgi) since DPAP3 is continuously expressed from early schizogony until egress. The facts that DPAP3 is active under mild acidic condition and that it is labelled by FY01 at pH 7.2 (Fig. 3) suggest that it might be able to cleave its natural substrate a neutral pH. However, we think it is unlikely that DPAP3 acts extracellularly on RBC surface proteins because co-culturing equal amounts of WT and DPAP3KO parasites did not rescued the DPAP3KO invasion defect (Fig. S7b).

We hypothesize that DPAP3 processes the N-terminus of adhesin proteins on the merozoite surface, either during traffic or after secretion, thus changing their affinity toward RBCM receptors. Most merozoite surface proteins are processed during traffic or after secretion, thus exposing one or multiple extracellular free N-termini that might be potential DPAP3 substrates^7,44^. These include most, if not all, type I transmembrane surface proteins, which expose their N-terminal ectodomain (EBAs, Rhs, etc), GPI-anchored proteins (MSP1, SUB2, P113), and surface proteins interacting with membrane-anchored protein (Rh5, Ripr, MSP6&7). To the best of our knowledge the specific functions of these N-terminal residues have not been studied, however the N-terminal region of some of these proteins has been shown to be key in mediating protein-protein interactions important for RBC invasion. These include interaction between GPI-anchored P113 and the N-terminal of Rh5^45^, between the N-terminal domain of Rh4 and host CR1^46^, or between the N-terminal of AMA1 and RON2. Also, conformational changes of the N-terminal tail of micronemal PfTRAP and TgMIC2 have been shown to be important for invasion of hepatocytes by *Plasmodium* sporozoites^47^, and for motility of the related Apicomplexan parasite *Toxoplasma gondii*^48^, respectively. Interestingly, *T. gondii* also expresses DPAPs in secretory organelles^49^, and a recent proteomic study has shown evidence that the N-terminus of certain secreted proteins, such as TgSUB1, TgMIC11, and TgSUB1, might be trimmed through the removal of dipeptides^50^. It is therefore possible that DPAP3-mediated trimming of the N-terminus of certain merozoite surface proteins might modulate the function of adhesin factors. Interestingly, during our invasion assays we observed a lot of variation in the effect of DPAP3 KO on RBC invasion, ranging from 25-75% inhibition (Fig. 8b). We think this variation might reflect heterogeneity among the different batches of blood used to perform invasion assays and suggests that DPAP3 activity might be important to recognize specific RBCM receptors that are differentially expressed in the human population.

Finally, our studies show that DPAP3 is an unusual papain-fold protease since removal of its prodomain is not required for activation. Most protease prodomains act as endogenous inhibitors and internal chaperones. Although the prodomain of DPAP3 might be required for proper folding of this large protein (941aa), we cannot discard the possibility that it might have other biochemical functions, such as recognizing substrates or binding to cofactors that modulate DPAP3 activity. Interestingly, the pro-form of *P. falciparum* DPAP1 has been shown to localize to the PV in mature schizonts^28^, and processing of recombinant DPAP1 ‘zymogen’ form with papain or trypsin only increases its activity 2-3 fold^30^, suggesting that similarly to DPAP3, its ‘zymogen’ form might be active and able to process the N-terminus of merozoite surface proteins. Thus, DPAP1 and DPAP3 might play redundant or complementary functions during RBC invasion, and further genetic and inhibitor studies are required to determine whether inhibitors with dual specificity would completely block RBC invasion.

## Methods

Synthesis of inhibitors, production of DPAP3 antibodies, and a list of primers (Table S2) and antibodies (Table S3) used in this study are shown in the supplementary methods.

### Design of expression constructs

Three synthetic genes codon-optimized for insect cells were synthesized by Genewiz and cloned into the puc57 vector backbone: puc57-rDPAP3-Nt, puc57-rDPAP3-Ct-wt, and puc-rDPAP3-Ct-mut. The first one codes for the N-terminal portion of DPAP3 (Met1-Asp469) and the other two for the C-terminal portion (Lys455-Stop941) containing the catalytic domain of DPAP3 and harboring either the catalytic cysteine Cys504 or the C504S inactivating mutation. All synthetic sequences contained a C-terminal His6-tag and were flanked with the BamHI and HindIII restriction sites at the 5’ and 3’ end, respectively. A ClaI restriction site is present in the 45bp overlapping sequence (Lys455-Asp469) between puc57-rDPAP3-Nt and puc57-rDPAP3-Ct-wt/mut. Digestion of these plasmids with BamHI, HindIII, and ClaI, followed by ligation of the N- and C-terminal products into the puc57 backbone yielded puc57-rDPAP3-WT and puc57-rDPAP3-MUT. The BamHI and HindIII restriction sites were used to clone full length *dpap3* into the pFastBacHT vector (Thermo Fisher Scientific) for expression of WT or MUT DPAP3 (pFB-rDPAP3wt and pFB-rDPAP3mut) in insect cells.

### Recombinant expression of DPAP3

DPAP3 was expressed in Sf9 insect cells using the baculovirus system. *E. coli* DH10Bac cells (Invitrogen) were transformed with pFB-rDPAP3wt and pFB-rDPAP3mut following the manufacturer recommendation. Baculovirus DNA was extracted using the BACMAX^TM^ DNA purification kit (Epicentre) and transfected into a 5mL culture of Sf9 cells (2×10^6^ cells/mL) using Cellfectin (Thermofisher). After 3 days, the culture supernatant containing baculovirus particles was collected (P1 stock). To increase the viral load of our stocks, 1mL of culture supernatant was serially passage twice into 25mL of Sf9 cultures at 2×10^6^ cell/mL for 3 days to obtain P2 and P3 viral stocks, which were stored at 4°C or frozen in liquid N_2_ in the presence of 10% glycerol. Insect cells were grown in SF-900-II serum free medium (Gibco) at 27°C under shaking conditions.

For rDPAP3 expression, Sf9 cells at 2×10^6^ cells/mL were infected with 0.4mL of P3 viral stock per liter of culture. The supernatant containing active rDPAP3 was collected 72h after infection, supplemented with protease inhibitors (1mM PMSF, 0.5mM EDTA, 1μM pepstatin, 1μM bestatin, and 10μM E64), and its pH adjusted by adding 50mM TrisHCl from a 1M solution at pH 8.2; 10% glycerol was added before storage at −80°C. (Note that despite being a general Cys protease inhibitor, E64 does not inhibit DPAP3.)

A three steps purification consisting of ion exchange, affinity, and size exclusion chromatography was used to purify rDPAP3. First, culture supernatant was passed 3 times through 0.05 volumes of Q-sepharose (GE Healthcare) pre-equilibrated with Buffer A (50mM Tris pH 8.2 containing the above-mentioned protease inhibitors), the resin washed with 5 volumes of Buffer A and 2.5 volumes of 50mM NaCl in Buffer A. Recombinant DPAP3 was eluted with 400mM NaCl in Buffer A. Fractions containing active DPAP3 (measured by VR-ACC turnover and FY01 labelling) were pooled, diluted 1:1 into Buffer B (100mM sodium acetate, 100mM NaCl, pH6, and protease inhibitors mentioned above), and passed through 0.05 volumes of Ni-NTA resin (Qiagen). The resin was washed with 10 volumes of Buffer B, and rDPAP3 eluted by lowering the pH of Buffer B to 5. Fractions containing active DPAP3 were pooled, concentrated using a Centricon Plus-70 Centrifugal Filter Unit (Millipore), loaded on a SuperdexTM 200 10/300 GL size exclusion column, and run on an AKTA FPLC with Buffer A. Active fractions were pooled, concentrated, and store at −80°C in the presence of 10% glycerol.

### DPAP3 activity assay using fluorogenic substrates or FY01

DPAP3 activity was measured either using the DPAP fluorogenic substrates VR-ACC^30^ or FR-βNA (Sigma), or with the FY01 activity-based probe. When using FY01, samples (intact parasites, parasite lysates, insect cell supernatant, or rDPAP3 purification fractions) were labelled with 1μM FY01 for 1h, boiled in loading buffer, run on a SDS-PAGE gel, and the fluorescence signal measured on a PharosFX (Biorad) flatbed fluorescence scanner^17^. To determine the potency and specificity of inhibitors against DPAPs and the falcipains, parasite lysates diluted in acetate buffer (50mM sodium acetate, 5mM MgCl_2_, 5mM DTT, pH 5.5) were pretreated for 30min with a dose response of inhibitor followed by FY01 labelling.

When using VR-ACC (10μM) or FR-βNA (100μM), substrate turnover was measure on a M5e Spectramax plate reader (λ_ex_=355nm/λ_em_=460nm or λ_ex_=315nm/λ_em_=430nm, respectively) in 50mM sodium acetate, 20mM NaCl, 5mM DTT, and 5mM MgCl_2_, pH5.5. The pH dependence of rDPAP3 was determine at 10uM VR-ACC using a 20mM sodium acetate, 20mM MES and 40mM TRIS triple buffer system containing 5mM DTT, 0.1% CHAPS, 20mM NaCl, and 5mM MgCl_2_.

### Design of targeting and complementation constructs

All constructs designed to integrate into the *dpap3* locus by single-crossover recombination (tagged or cKO lines) contained either a 1065bp C-terminal homology region fused to GFP, or a 1210bp homology region upstream of the catalytic Cys504 (Asp39-Glu392) fused to a recodonized C-terminal region (Lys393-Stop941) tagged with mCherry or HA_3_. The recodonized region contained either WT Cys504, or the C504S mutation. The construct designed to generate the DPAP3-HA, DPAP3-mCh, and DPAP3-GFP tagged lines are shown in Fig. S8a-b. Briefly, the *dpap3* C-terminal homology region was inserted into the pPM2GT plasmid to generate pPM2GT-DPAP3Ct-GFP. The N-terminal region of *dpap3* in the pFB-rDPAP3-wt/mut vector was replaced with the homology region to generate pFB-chDPAP3-wt/mut. The *dpap3* ORF was then introduced into the pHH1-SERA5ΔCt-HA—obtained after removal of the C-terminal part of SERA5 in the pHH1-SERA5-loxP-DS_PbDT3’^31^—resulting in plasmids pHH1-chDPAP3-wt/mut-HA, which harbor a C-terminal HA_3_ tag and a *loxP* site downstream of the Pb3’ UTR. The HA_3_ sequence of pHH1-SERA5ΔCt-HA was replace with mCh—amplified from the pREST-B plasmid^51^—to generate the pHH1-SERA5ΔCt-mCh, and the ORF of WT or MUT DPAP3 introduced into this plasmid to generate the pHH1-chDPAP3-wt/mut-mCh constructs.

To conditionally truncate *dpap3* we used two different approaches. The first approach introduced a *loxP* site within the ORF of *dpap3*, in an Asn-rich region (Asn414-Asn444) upstream of the catalytic domain, resulting in replacement of Asn430-Asp434 with a *loxP* coding peptide (ITSYSIHYTKLFTG). To make the pHH1-chDPAP3_loxP-mCh construct (Fig. S8c), the N- and C-terminal portions of DPAP3 were amplified from the pHH1-chDPAP3-wt-mCh, which contains a 3’UTR *LoxP* site, and ligated into the plasmid backbone. A *LoxP* site was introduced in the backward primer used to amplified the N-terminal region. The second approach inserted a *loxPint*^34^ between the homology and recodonized regions. A synthetic 1600bp sequence (GeneWiz®) containing the *loxPint* fragment flanked by targeting sequences was introduced into construct pHH1-chDPAP3-wt-mCh to generate the pHH1-chDPAP3_loxPint-mCh plasmid.

Finally, for episomal complementation of the cKO lines, plasmids pHH1-chDPAP3-wt/mut-HA (Fig. S8a) were modified in order to express full-length WT or MUT DPAP3 under the control of the *dpap3* or *ama1* promoters. Firstly, to select for parasites containing the complementation plasmids after transfection, the puromycin N-acetyltransferase (pac) gene, which confers resistance to puromycin, was amplified from mPAC-TK (a kind gift of Alex Maier) and subsequently ligated into pHH1-chDPAP3-wt/mut-HA plasmids. The homology region of this plasmid was replaced with a recodonized N-terminal *dpap3* amplified from puc57-rDPAP3-Nt, resulting in pHH1-rDPAP3-wt/mut-HA plasmids (Fig. S8d). The *dpap3* and *ama1* promoters (970 and 1456bp upstream of the start codon, respectively) were amplified from genomic DNA and ligated into these plasmids to generate the pHH1-*dpap3-*rDPAP3-wt/mut-HA, and pHH1-*ama1-*rDPAP3-wt/mut-HA complementation constructs.

All final construct sequences were verified by nucleotide sequencing on both strands. Primers for PCR amplification and restriction sites used to generate these plasmids are indicated in Fig. S8 and listed in Table S2.

### Generation and maintenance of transgenic parasite lines

All constructs were transfected at schizont stage using a 4D-Nucleofector electroporator (Lonza) as previously described^12^. DPAP3-tagged and DPAP3-cKO lines were obtained through multiple on and off drug selection cycles with WR99210 (Jacobus Pharmaceuticals) and cloned by limited dilution as previously described^52^. To generate the DPAP3-GFP, DPAP3-HA and DPAP3-mCh lines, pPM2GT-DPAP3Ct-GFP, pHH1-chDPAP3-wt-HA and pHH1-chDPAP3-wt-mCh were transfected into *P. falciparum* 3D7 parasites. Plasmid pHH1-chDPAP3-mut-mCh was transfected multiple times into 3D7 in an attempt to swap the endogenous catalytic domain of DPAP3 with one containing the inactivating C504S mutation, but no integration was observed even after five drug cycles. The A1cKO and F3cKO & F8cKO lines were obtained after transfection of pHH1-chDPAP3_loxP-mCh and pHH1-chDPAP3_loxPint-mCh, respectively, into *P. falciparum* 1G5 parasites that endogenously expressed DiCre. Finally, the E7ctr line containing only the 3’UTR *loxP* site was generated by transfecting 1G5 parasites with pHH1-chDPAP3-wt-mCh. Complementation lines were obtained after transfection of A1cKO or F8cKO with pHH1-*dpap3-*rDPAP3-wt-HA, pHH1-*dpap3-*rDPAP3-mut-HA, pHH1-*ama1-*rDPAP3-wt-HA, or pHH1-*ama1-*rDPAP3-wt-HA and selection with WR99210 and puromycin. All parasite lines were maintained in RPMI 1640 medium with Albumax (Invitrogen) containing WR99210 (plus puromycin for the complementation lines) and synchronized using standard procedures^53^.

To conditionally truncate the catalytic domain of DPAP3 tightly synchronized ring-stage parasites were treated for 3-4h with 100nM RAP (Sigma) or DMSO at 37°C, washed with RPMI, and returned to culture^31^. Schizonts purified at the end of the cycle were used to determine the excision efficiency at the DNA (PCR) or protein (IFA, WB) level.

### Staining and microscopy

Thin films of *P. falciparum* cultures were air-dried, fixed in 4% (w/v) formaldehyde (PFA) for 20min (Agar Scientific Ltd.), permeabilized for 10min in 0.1% (w/v) Triton X100 and blocked overnight in 3% (w/v) bovine serum albumin (BSA) or 10% (w/v) goat serum (Invitrogen) in PBS. Slides were probed with monoclonal antibodies or polyclonal sera as described previously^54^ (See Table S3 for antibodies used in this study), subsequently stained with Alexa488-, Alexa594-, Alexa647-labelled secondary antibodies (Molecular Probes) and DAPI (4,6-diamidino-2-phenylindole), and mounted in ProLong Gold Antifade (Molecular Probes). Images were collected using AxioVision 3.1 software on an Axioplan 2 Imaging system (Zeiss) using a Plan-APOCHROMAT 100x/1.4 oil immersion objective or LAS AF software on an SP5 confocal laser scanning microscope (Leica) using a HCX PL APO lamda blue 63x/ 1.4 oil immersion objective.

Super-resolution microscopy was performed using a DeltaVision OMX 3D structured illumination (3D-SIM) microscope (Applied Precision). Images were analyzed with ImageJ (NIH), Adobe Photoshop CS4 (Adobe Systems) and Imaris x64 9.0.0 (Bitplane) software.

### Immunoelectron microscopy

For IEM, mature schizonts from the DPAP3-GFP and 3D7 control lines were concentrated using a magnetic activated cell sorting (MACS) LD separation column (Miltenyi Biotec). Briefly, iRBCs were loaded onto an LD column attached to Midi MACS pre-equilibrated with media. The column was washed twice with media and schizonts eluted with media after detaching the column from the magnet. Parasites were then fixed in 4% paraformaldehyde/0.1% glutaraldehyde (Polysciences) in 100 mM PIPES and 0.5mM MgCl_2_, pH 7.2, for 1h at 4°C. Samples were embedded in 10% gelatine and infiltrated overnight with 2.3M sucrose/20% polyvinyl pyrrolidone in PIPES/MgCl_2_ at 4°C. Samples were trimmed, frozen in liquid nitrogen, and sectioned with a Leica Ultracut UCT cryo-ultramicrotome (Leica Microsystems). Sections of 70nm were blocked with 5% FBS and 5% NGS for 30min and subsequently incubated with rabbit anti-GFP antibody 6556 (Abcam) at 1:750 overnight at 4°C. Colloidal gold conjugated anti rabbit (12 nm) IgG (Jack Imm Res Lab) was used as secondary antibody.

### Plaque assays

Plaque assays were performed as previously described^16^. Briefly, the 60 internal wells of a flat-bottom 96 well plates were filled with 200µL of DMSO- or RAP-treated parasite culture at 10 iRBCs/well and 0.75% hematocrit, incubated for 12-14 days at 37°C, and the number of microscopic plaques counted using an inverted microscope.

### Flow cytometry-based replication, time course, and invasion assays

For all FACS-based assays, samples were fixed for 1h at RT with 4% PFA and 0.02% glutaraldehyde, washed with PBS, and stored at 4°C. Samples were stained with SYBR Green (1:5000) or Hoechst (2μg/mL), run on a FACScalibur or FortessaX20 flow cytometers (Becton-Dickinson Bioscience), and the data analyzed with CellQuest Pro or FlowJo.

For replication assays, cultures at 0.1% parasitemia (ring stage) and 2% hematocrit were grown for 4 cycles after DMSO or RAP treatment. Aliquots were fixed every 48h at trophozoites stage, stained with SYBR Green, and parasitemia quantified by flow cytometry. To avoid parasite overgrowth, cultures were diluted 10-times whenever they reached 5% parasitemia. The cumulative percentage parasitemia (CP) over 4 cycles was fitted to an exponential growth model: CP=P_t0_.MR^N^, where P_t0_ is the initial parasitemia, MR the multiplication rate per cycle, and N the number of cycles after treatment.

To look at the effect of DPAP3 truncation on the full erythrocytic cycle, A1cKO parasites were synchronized within a 2h window, treated with DMSO or RAP for 3h, and put back in culture for 76h. Sample were collected every 2-4h, fixed, stained with Hoechst, and analyzed by flow cytometry. DNA levels were quantified as the median fluorescence signal of iRBC divided by the background signal for uninfected RBCs (uRBCs).

For standard invasion assays, schizonts purified from DMSO or RAP treated cultures were incubated with fresh RBCs for 8-14h, fixed, and stained with Hoechst. The population of uRBC, rings and schizonts was quantified based on DNA content. The invasion rate was determined as the ratio between the final population of rings and the initial population of schizonts. Invasion time courses were performed by arresting purified schizonts with 1μM C2 for 4h, washing twice with warm media, and culturing with fresh RBCs under shaking conditions. Samples were collected at different time points, fixed, and split into two aliquots: One was stained with Hoechst and WGA-Alexa647 and run on a flow cytometer at a high forward scattering voltage in order to detect free merozoites. The populations of uRBC, free merozoites, rings, and schizonts were quantified with FlowJo. The other aliquot was blocked with 3% BSA in PBS overnight at 4°C, and stained without permeabilization with the MSP1 monoclonal antibody 89.1 (1:100), and subsequently with anti-mouse Alexa488 (1:3000). The population of uRBC, schizonts, rings, and merozoites attached to the RBC surface were quantified using FlowJo.

### Live microscopy imaging of egress and DPAP3 secretion

Time lapse video microscopy of egress was performed as previously described^9^. Briefly tightly synchronized schizonts were percoll-enriched and arrested with 1µM C2 for 4h. After C2 wash out, DIC and mCherry images were collected every 5 and 25s, respectively, for 30min using a Nikon Eclipse Ni-E wide field microscope fitted with a Hamamatsu C11440 digital camera and a Nikon N Plan Apo λ 100x/1.45NA oil immersion objective. For each experiment, videos of the RAP-and DMSO-treated parasites were taken alternately to ensure that differences in the rate of egress were not a result of variation in the maturity of the parasite populations. The images were then annotated using Axiovision 3.1 software and exported as AVI movie or TIFF files. Individual egress events were annotated by detailed visual analysis of the movies, and the delay to the time of egress was recorded for each schizont for subsequent statistical analysis. Mean fluorescence intensity values of individual mCherry-expressing schizonts right before and after PVM breakdown were determined from exported raw image files (TIFF format) as described previously^9^ and using the elliptical selection tool and ‘Histogram’ options of ImageJ/Fiji V1.0. DPAP3KO parasites were analyzed to determine the residual background fluorescence derived from the hemozoin.

### Western blots

Saponin pellets of parasites from different erythrocytic stages as well as samples from culture supernatant harvested during egress and invasion were syringe filtered (Minisart, 0,2 µm, Sartorius), boiled in SDS-PAGE loading buffer under reducing conditions, run on a SDS-PAGE gel, and transferred to Hybond-C extra nitrocellulose membranes (GE Healthcare). The membranes were blocked with 5% (w/v) nonfat milk in PBS, probed with monoclonal or polyclonal antibodies (see Table S3 for antibodies used in this study), and followed by application of horseradish peroxidase-conjugated secondary antibodies (Pierce). The signal was detected using SuperSignal West Pico chemiluminescent substrate (Thermo Scientific) and a ChemiDoc MP imager (BioRad).

### Diagnostic PCR

PCR was performed using GoTag (Promega), Advantage 2 (Clontech), or Q5 High-Fidelity (NEB) polymerases. Diagnostic PCR to detect integration of targeting constructs was performed using extracted genomic DNA as template. Primer pairs specific for detection of integration, namely P21 and P22 for integration of pHH1-chDPAP3-mCh and pHH1-chDPAP3-HA, and II-inte_F and II-wt_R for integration of pPM2GT-DPAP3Ct-GFP, were designed such that the forward primer hybridized in a genomic region upstream of the plasmid homology region, and the second in a region unique to the introduced plasmid. Primer pairs P21 and P23 were designed to detect presence of the unmodified *dpap3* locus. To assess whether *dpap3-mCh* had been excised after RAP treatment, diagnostic PCR was performed using extracted genomic DNA as template. Primers P24 and M13 were used to detect non-excision *dpap3* at the genomic locus and hybridize upstream and downstream of the second *loxP* site, respectively. Primers P25 and SP6 were used to detect presence of excision and hybridize upstream and downstream of the first and second *loxP* sites, respectively.

## Acknowledgements

We would like to thank Prof. Michael Blackman and his group for their support, assistance and suggestions about this work, and for providing antibodies, the 1G5 parasite cell line, and plasmids to generate several constructs. We also thank Dr Anthony Holder, Dr. Jean-Francois Dubrenetz, Dr. Alan Thomas, and MR4 for providing several antibodies, Dr Alex Maier for providing the mPAC-TK plasmid, and Dr Christiaan van Ooij for providing primers. We also thank Prof. Matthew Bogyo for providing the FY01 probe and MRCT for providing compound 2. We would also like to thank Prof. Klemba at Virginia Tech for its contribution in generating the DPAP3-GFP line, and Dr Wandy Beatty at Washington University, St Louis, for immunoelectron microscopy support. Finally, we thank the Wellcome Trust and Royal Society for funding this research and E.D. and C.L. through the Sir Henry Dale Fellowship 099950, Erasmus for funding L.E.V., and the National Institute for Medical Research for funding M.S.Y.T.

## Author Contributions

C.L. performed most of the experimental work and contributed to the writing of the manuscript. M.S.Y.T. and L.E.V. expressed, purified, and characterized recombinant DPAP3, and M.S.Y.T. also conducted experiments to study DPAP3 processing and secretion. I.R. generated the DPAP3-GFP lines and performed the IEM studies under the supervision of D.A.G. M.I.S. synthesized SAK1, *L*-WSAK, and *D-*WSAK. E.D. performed experiments, supervised the work, and wrote the manuscript.

